# In Vitro Efficacy of Next Generation Dihydrotriazines and Biguanides Against Babesiosis and Malaria Parasites

**DOI:** 10.1101/2024.03.20.585986

**Authors:** Pratap Vydyam, Meenal Chand, Shalev Gihaz, Isaline Renard, Gavin D. Heffernan, Laura R. Jacobus, David P. Jacobus, Kurt W. Saionz, Raju Shah, Hong-Ming Shieh, Jacek Terpinski, Wenyi Zhao, Emmanuel Cornillot, Choukri Ben Mamoun

## Abstract

*Babesia* and *Plasmodium* pathogens, the causative agents of babesiosis and malaria, are vector-borne intraerythrocytic protozoan parasites, posing significant threats to both human and animal health. The widespread resistance exhibited by these pathogens to various classes of antiparasitic drugs underscores the need for the development of novel and more effective therapeutics strategies. Antifolates have long been recognized as attractive antiparasitic drugs as they target the folate pathway, which is essential for the biosynthesis of purines and pyrimidines, and thus are vital for the survival and proliferation of protozoan parasites. More efficacious and safer analogs within this class are needed to overcome challenges due to resistance to commonly used antifolates such as the aminopyrimidine, pyrimethamine, and to address liabilities associated with the dihydrotriazines, WR99210 and JPC-2067. Here we utilized an in vitro culture condition suitable for continuous propagation of *B. duncani, B. divergens, B. MO1* and *P. falciparum* in human erythrocytes to screen a library of 51 dihydrotriazines and 28 biguanides for their efficacy in vitro and to compare their potency and therapeutic indices across different species and isolates. We identified nine analogs that inhibit the growth of all species, including the *P. falciparum* pyrimethamine-resistant strain HB3 with IC_50_ values below 10 nM and demonstrated excellent therapeutic indices. These compounds hold substantial promise as lead antifolates for further development as broad-spectrum antiparasitic drugs.

## Introduction

The hemoparasites *Plasmodium* and *Babesia* invade host red blood cells where they develop and multiply to cause the pathological symptoms associated with human malaria and babesiosis, respectively. Whereas *Plasmodium* species are transmitted by mosquitoes, *Babesia* parasites are primarily transmitted by ticks. The 2023 WHO report indicated that, in 2022, malaria was responsible for 249 million clinical cases and 608,000 deaths worldwide, the majority of which were caused by *P. falciparum* [1]. Human babesiosis has also a worldwide distribution with most cases reported in the United States where the disease is considered endemic and represents a significant concern to the blood supply [1–3]. Several *Babesia* species have been shown to cause human disease [3, 4]. These include *B. microti, B. duncani and Babesia MO1* in the US, and *B. divergens* in Europe [5]. The number of reported cases of human babesiosis has increased significantly since 2011 when it became a nationally notifiable disease [2, 5, 6]. The CDC recently reported that over 14,000 cases of human babesiosis were reported between 2011-2018 in the United States [2]. Although transmission of *Babesia* parasites is primarily via the tick bite, other transmission modes have also been reported with blood transfusion being the most common [7, 8]. The disease manifestations range from asymptomatic to severe, with typical clinical symptoms including chills, fever, and sweats. For people with weak immune systems (such as cancer, lymphoma, and AIDS), immunological complications, and immune therapies, the disease can be more severe and sometimes lethal [9]. The current CDC-recommended treatment protocol for human babesiosis includes combinatorial therapy of the antimalarials azithromycin with atovaquone for mild babesiosis and clindamycin with quinine for severe babesiosis [2, 10]. However, these treatments are not very effective and are often associated with adverse events [11–13]. Consequently, persistent *Babesia* infections and recrudescence can lead to prolonged treatment [14, 15] and emergence of mutant parasites, which are resistant to therapies [16, 17]. Similar mutations were also identified following infection of mice with *B. microti* or *B. duncani* [18, 19] or selected following drug pressure in vitro [20, 21]. Altogether, these findings highlight the need for more effective therapies for the treatment of human babesiosis.

Because folate metabolism has long been known to be essential for the development and survival of parasites, targeting this process by inhibiting specific enzymatic steps in biosynthesis of folates has long been a major focus of the antiparasitic drug development efforts for both prophylaxis and treatment [22, 23]. Folate cofactors are essential in several metabolic processes including the production of purine and pyrimidine for DNA replication, and amino acid metabolism. First generation antifolates have been shown to act as competitive inhibitors of bifunctional enzyme dihydrofolate reductase thymidylate synthases (DHFR-TS) from protozoan parasites, and have been used as antiparasitic drugs against *Plasmodium* and *Toxoplasma* species since World War II [22, 24, 25]. The use of antifolate-based treatments is often applied when parasites become resistant to other first-line antiparasitic treatments. One such antifolates is pyrimethamine (PYM) (2,4-Diamino-5-(4-chlorophenyl)-6-ethylpyrimidine), which selectively inhibits the DHFR-TS enzymes of protozoan parasites [26], and is commonly used in combination with other inhibitors of other steps in the folate metabolic pathway, including sulfadoxine (drug combination known as Fansidar) and dapsone (drug combination known as Maloprim) for maximum efficacy [27, 28].

A major challenge to the efficacy of antifolate-based drug therapy arose with the emergence of antifolate-resistant isolates [29, 30]. Parasite lines with point mutations in the DHFR-TS enzyme showed reduced susceptibility to treatment with a combination of sulfonamide and pyrimethamine [31–33]. The prevalent mutations in *Pf*DHFR-TS associated with resistance to pyrimethamine include N51I, C59R, S108N, and I164L. These mutations result in decreased binding affinity of antifolates for the DHFR-TS enzyme due to spatial restrictions imposed by the drug, leading to a reduction in binding efficacy [34, 35]. Field and in vitro studies have shown that S108N is the most crucial substitution in *Pf*DHFR-TS responsible for resistance to this drug [22, 32, 33, 36, 37].

The small molecule WR99210 is a second generation antimalarial triazine derivative developed in the late 1960s with improved efficacy against pyrimethamine- and chloroquine-resistant parasites [38, 39]. A mutagenesis library study of the DHFR-TS gene in *E. coli* uncovered a reverse mutation (N108S) that augments binding affinity of WR99210, leading to heightened sensitivity of parasites to these compounds [34, 40]. However, WR99210 was found to have poor bioavailability and to cause gastrointestinal complications in healthy subjects resulting in its withdrawal from further clinical evaluation [41, 42]. Third generation biguanide prodrugs has been reported to show improved gastrointestinal tolerability and antiparasitic activity in malaria. [39, 41]. One of these, JPC-2067, showed promise when tested against an array of pathogens including *P. falciparum, T. gondii, Mycobacterium tuberculosis,* and *Nocardia species* [43–45]. Unlike their efficacy against *Plasmodium* and *Toxoplasma* species, pyrimethamine, sulfamethoxazole, trimethoprim, and dapsone monotherapy or the combination of Pyrimethamine-sulfadoxine have been shown to have little to no activity against *Babesia microti* infection in Mongolian jirds [46]. A moderate to effective inhibition of *in vitro* parasite growth of *Babesia gibsoni* and *Babesia bovis* was observed with trimethoprim, methotrexate, and pyrimethamine [47, 48].

Recent advances in understanding the genomic composition of various *Babesia* species that infect humans have helped identify potential targets for more effective anti-babesial drugs. Anti-babesial drug discovery was further accelerated following the development of the *B. duncani* model of infection, which combines both a continuous *in vitro* culture of the parasite in human red blood cells and infection in C3H/HeJ mice [49–51]. Interestingly, previous studies have shown moderate efficacy of pyrimethamine against *B. duncani* with an IC_50_ value of 940 nM, whereas WR99210 is highly potent against the parasite with an IC_50_ value of 0.5 nM [52].

In order to search for potent but safer analogs of WR99210, we standardized the in vitro culture systems of several intraerythrocytic parasites in human erythrocytes in order to compare drug efficacy across different species and isolates. Using this strategy, we screened a library of 79 compounds (51 dihydrotriazines and 28 biguanides), analogs of WR99210, against *B. duncani*, *B. divergens*, *B. MO1* (*B. divergens*-like species) and *P. falciparum* drug-sensitive and - resistant strains. Nine analogs with IC_50_ values <10 nM against all babesiosis and malaria parasites were identified and are promising early candidates for further development as broad-spectrum antiparasitic drugs.

## Results

### *Babesia* DHFR-TS enzymes are suitable targets for antibabesial drug discovery

The availability of annotated proteomes from several human *Babesia* parasites, including *B. microti, B. duncani, B. divergens* and *B.sp MO1*, has enabled the analysis of the primary structure of their DHFR-TS enzymes for predicting the susceptibility of these organisms to antifolates (**Fig. 1A**). In contrast to the *P. falciparum* enzyme, which comprises 608 amino acids, the DHFR-TS enzymes of *Babesia* species are comparatively smaller, ranging in size from 502 to 514 amino acids. This size difference primarily arises from insertions of 94 to 106 amino acids in the malarial enzymes (**Fig. 1B and Fig. S1**). Comparative analysis of amino acid identities and similarities within the DHFR domain revealed that the DHFR enzymes of *Babesia* species share the highest similarity with those of *P. falciparum* [52] (**Fig. 1B and Fig. S1**). In particular, the S108 residue associated with susceptibility to pyrimethamine in *P. falciparum* and other apicomplexan parasites corresponds to threonine in *B. microti*, *B. duncani*, *B. divergens, B. MO1* and *B. bovis* (**Fig. 1B**). Phylogenetic analyses conducted on the complete protein sequences of DHFR-TS enzymes from 52 apicomplexan parasites revealed a consistent alignment with the evolutionary trajectory of species within the *Apicomplexa* phylum. Notably, *B. microti* DHFR-TS appears as an out-group relative to other Piroplasmida (see **Fig. 1C**). Examination of the DHFR and TS moieties across various apicomplexan lineages in this study revealed nearly identical evolutionary patterns. Interestingly, while no insertions or deletions (indels) were detected in the TS moieties, they were observed in the DHFR moieties. Furthermore, analysis of the linker region between the DHFR and TS domains (see **Fig. S2**) categorized them into long or short types, with the shortest linker being longer than those found in other protozoan parasites, such as kinetoplastidae [O’Neil, 2003 #78]. Taken together, these findings strongly indicate a close evolutionary relationship among DHFR-TS enzymes in *Apicomplexa*, encompassing species like *B. duncani*, *B. microti*, and *Plasmodium* spp.

**Figure 1.**
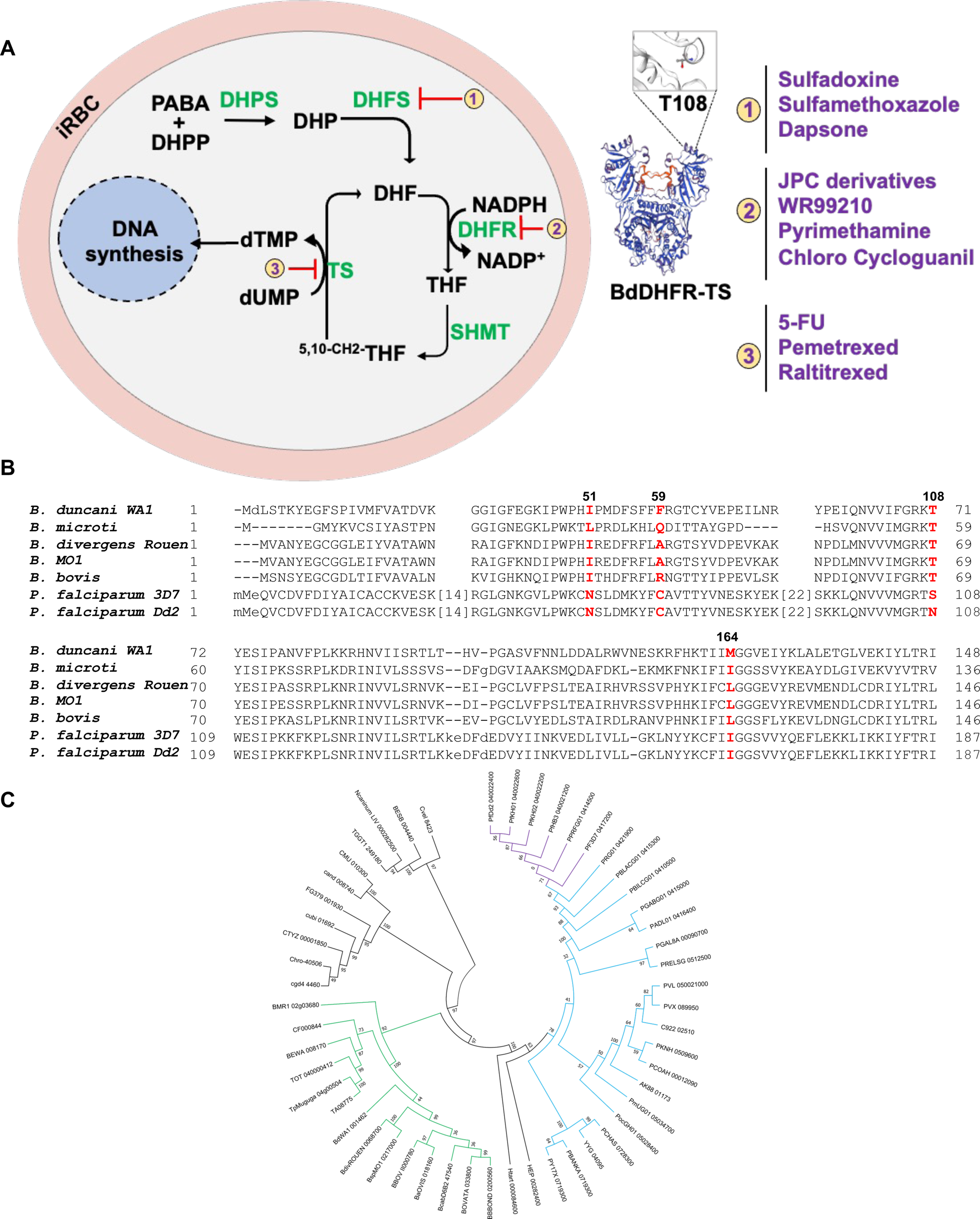
Conservation of the folate metabolic pathway among *Babesia* and *Plasmodium* parasites. **(A)** A schematic representation of folate metabolism in *Babesia* and *Plasmodium* parasites. The main enzymes (green color) targeted by antifolates (red color) are highlighted. Abbreviations (Abbs.): iRBC = infected red blood cell, DHP = dihydropteroate, DHPP = dihydropteroate pyrophosphate, DHPS = dihydropteroate synthase, DHF = dihydrofolate, DHFR = dihydrofolate reductase, PABA = p-aminobenzoic acid, SHMT = Serine hydroxy methyltransferase, THF = tetrahydrofolate, and TS = thymidylate synthase. **B)** Alignment of a region of DHFR-TS enzymes from various *Babesia* species as well as pyrimethamine-sensitive (3D7) and -resistant (Dd2) *P. falciparum* strains. Residues known to be associated with susceptibility or tolerance to antifolates are marked in red. **C)** Phylogenetic analysis of *DHFR-TS* enzymes from piroplasmids (tick-transmitted intraerythrocytic parasites) and various *Plasmodium* species. The tree was build using Phylogeny.fr pipeline combining Mafft, BMGE and FastTree softwares. An advanced option was used to generate 1000 boostraps providing branch supports. Clade supporting piroplasmida sequences is shown in green; *Plasmodium* sequences are in blue, with branches of *P. falciparum* sequences represented in violet.

### JPC-2067 inhibits parasite DHFR activity and the development of *B. duncani* in human erythrocytes

The in vitro antibabesial activity of both JPC-2067 and its pro-drug JPC-2056 (analogs of WR99210) was assessed against *B. duncani* using a continuous *in vitro* culture system in human erythrocytes. Both compounds exhibited inhibitory activity at the nanomolar level with IC_50_ values of 869.4 ± 0.3 nM and 8.9 ± 1.9 nM for JPC-2056 and JPC-2067, respectively (**Fig. 2A and 2B**). We further examined the cytotoxicity profile of both compounds against four human cell lines: HeLa, HCT-116, HepG2 and HEK-293, but no discernible effect was observed up to 900 µM. The calculated therapeutic indices ranged were notably high, exceeding 110 for JPC-2056 and more than 15000 for JPC-2067 (**Table 1**).

**Figure 2.**
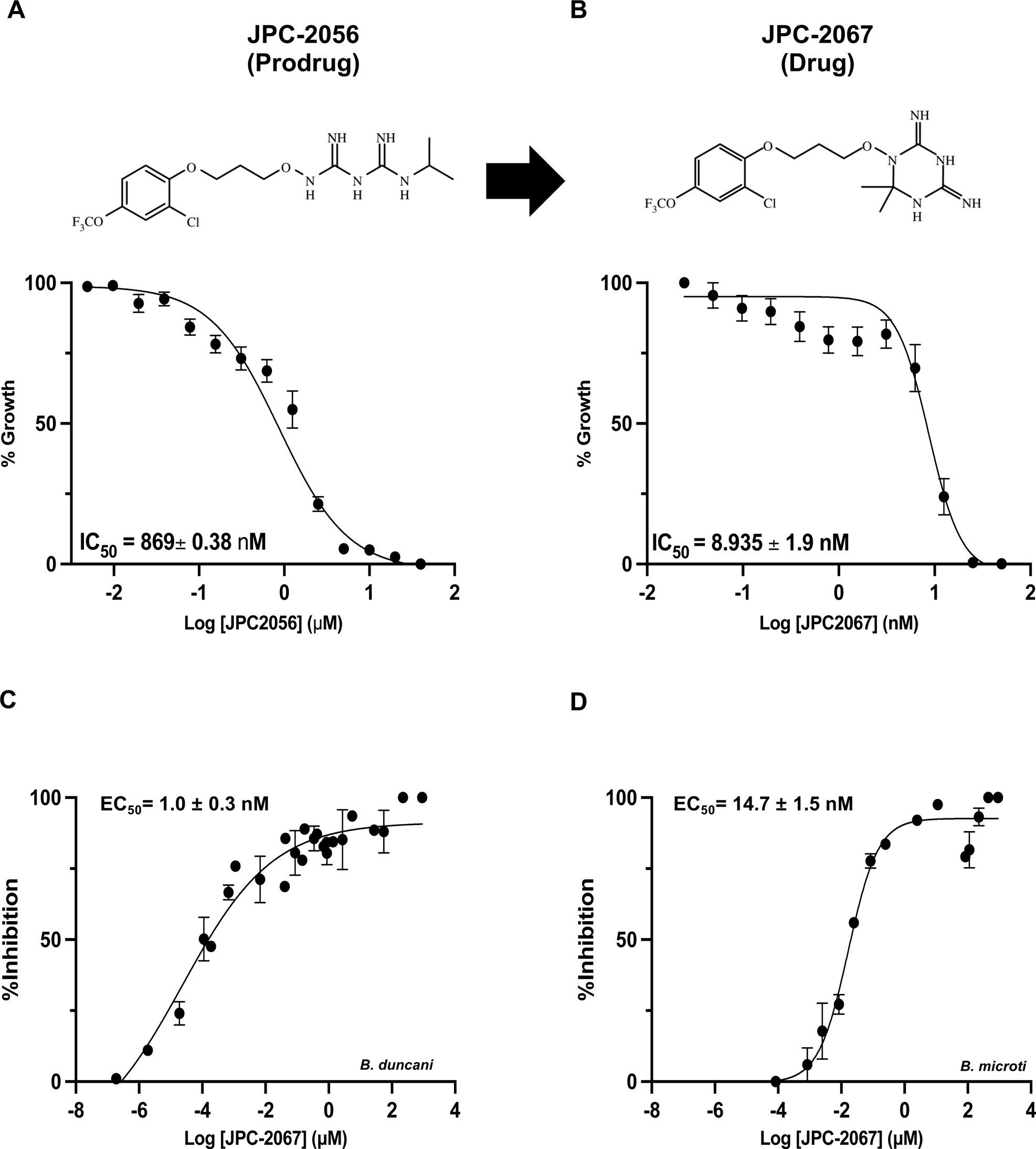
Inhibition of *B. duncani* DHFR-TS by JPC-2067 and JPC-2056. **(A and B)** Chemical structures and inhibition of *B. duncani* intraerythrocytic development by the prodrug, JPC-2056 (**A**), and its active drug JPC-2067 **(B)**. The data represent the mean of 2 independent experiments each in triplicate. Values represent mean ±SD. **C-D)** A dose-dependent inhibition of purified the DHFR activity of the *B. duncani* (**C**) and *B. microti* (**D**) DFR-TS enzymes by JPC-2067. EC_50_ values were calculated from the inhibition curves and represent mean ± SD from three independent experiments, each performed in triplicate.

**Table 1.** Cytotoxicity and therapeutic indices of JPC-2056 and its active metabolite JPC-2067.

| **Drug** | **IC<sub>50</sub>**<br>***B. duncani*** | **IC<sub>50</sub>**<br>**HeLa** | **IC<sub>50</sub>**<br>**HepG2** | **IC<sub>50</sub>**<br>**HEK293** | **IC<sub>50</sub>**<br>**HCT** | **Therapeutic**<br>**Index** |
| --- | --- | --- | --- | --- | --- | --- |
| **JPC2067** | 8.93 nM | >0.9 mM | 1.15 mM | 1 mM | 1 mM | >15000 |
| **JPC2056** | 869 nM | > 0.9 mM | 1 mM | 1 mM | 1 mM | >110 |

Recent studies have shown that the antifolate compound WR99210 inhibits the activity of DHFR-TS enzymes from *B. microti* and *B. duncani* [52]. Therefore, we examined the inhibitory activity of JPC-2067 against the *Babesia* enzymes. DHFR-TS activity measurement was performed in the absence or presence of rising concentrations of the compound. Dihydrofolate reductase (DHFR) catalyzes the NADPH-dependent reduction of dihydrofolate (DHF) to tetrahydrofolate (THF). Our studies revealed that JPC-2067 inhibit the enzymes with EC_50_ values of 1.0 nM and 14.7 nM for *B. duncani* and *B. microti*, respectively (**Fig. 2C and 2D**). In contrast, the prodrug JPC-2056 had no discernible effect on the *Babesia* enzymes (results not shown).

### In vivo efficacy of JPC-2056 in animal models of babesiosis

The finding that JPC-2067 is effective against *B. duncani* in vitro led us to investigate the activity of its prodrug JPC-2056 in C3H-HeJ mice. In this model, the mice infected with *B. duncani* parasites exhibit 100% mortality within 6-13 days post-infection [53]. To evaluate the efficacy of JPC-2056, we administered the drug orally continuously for ten days (DPI-1 to 10) to female mice infected with low doses (1×10^4^ iRBCs) of parasites. The group of mice treated with the vehicle (PEG-400) began showing signs of parasite infection within 5 days and reached peak parasitemia within ten days post-infection, with all animals succumbing to infection (**Fig. 3A and C**). Surprisingly, mice infected with parasites and subsequently treated with 30mg/kg of JPC-2056 displayed a slight delay in parasite appearance, eventually reaching peak parasitemia (>5%) by 12 days post-infection, and all animals succumbed to lethal *B. duncani* infection on day thirteen (**Fig. 3A**). Although JPC-2056 did not completely eliminate parasites from the mice, it did result in a one-day delay in parasite emergence and a significant difference (p=0.0003) in peak parasitemia were observed on the tenth day post-infection (**Fig. 3B**). Regardless of treatment, all mice groups developed disease symptoms and succumbed to *B. duncani* infection by day ten or thirteen post-infection (**Fig. 3C**).

**Figure 3.**
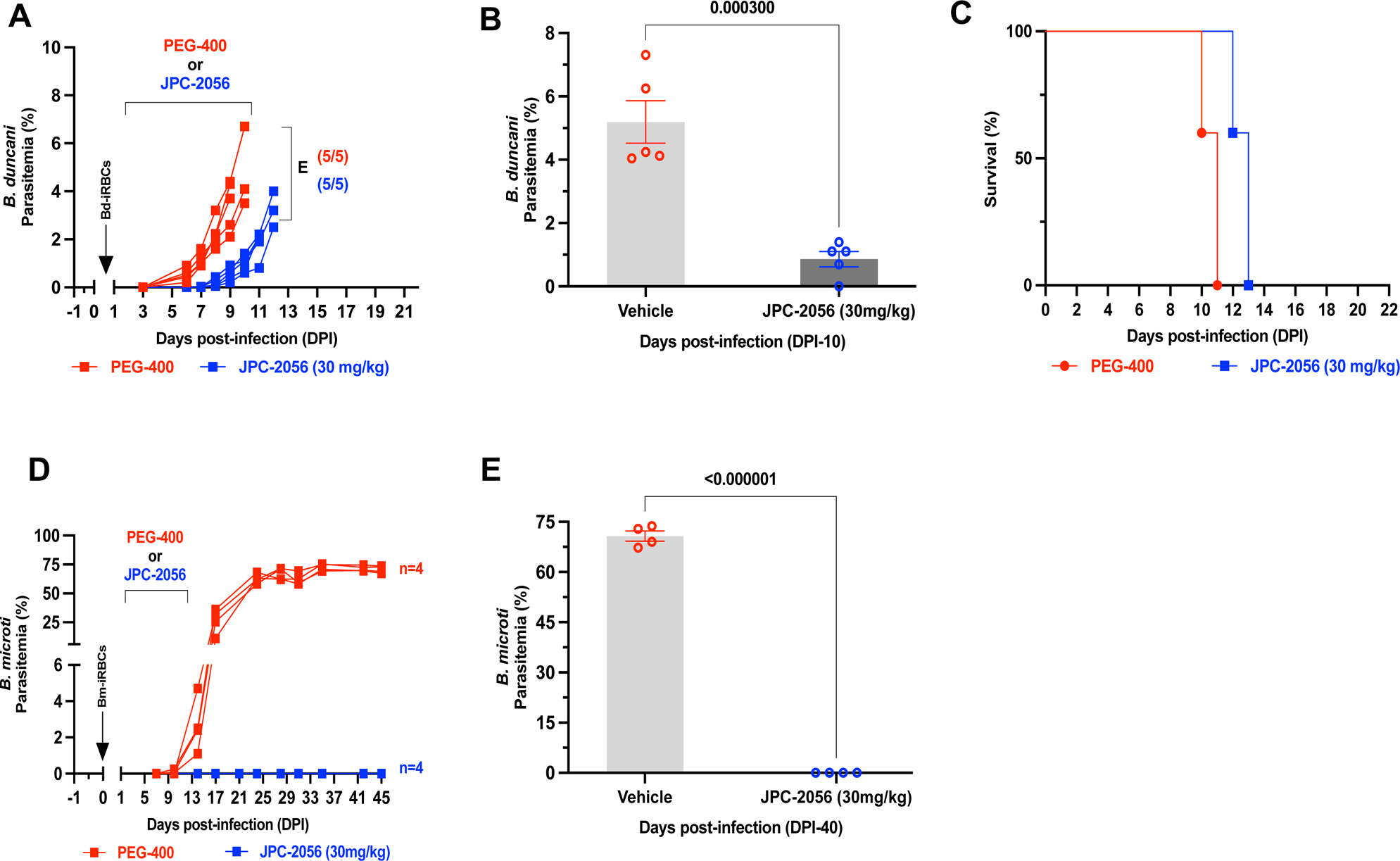
In vivo efficacy of JPC-2057 in mouse models of human babesiosis. **A)** Parasitemia in female C3H/HeJ mice (n=5) infected (intravenously) with 1×10^4^ *B. duncani*-iRBCs, which cause lethal infection in these mice, and treated with either vehicle (PEG-400) (red) or JPC-2056 (30 mg/kg, blue lines) administered via oral gavage once a day for 10 days (DPI 1-10). Treated mice were monitored daily and were euthanized when treatment was deemed not to be effective. Each data point depicts parasitemia measured via inspection of Giemsa-stained blood smears on the specified day. **B)** Bar diagram depicting % parasitemia in the vehicle and JPC-2056 groups on the last day of treatment (DPI10). Data are presented as mean ± SEM (n = 5 mice per group). **C)** Kaplan-Meyer curve depicting survival of *B. duncani-*infected female C3H/HeJ mice followed by treatment with vehicle and JPC-2056.**D)** Parasitemia in female CB17-SCID mice (n=5) infected with 10^4^ *B. microti* infected RBCs (LabS1 strain) and treated with either vehicle (PEG-400, red lines) or JPC-2056 (blue lines) at 30 mg/kg once a day for 10 days. **E)** Bar diagram % parasitemia after 40 days post-infection in the vehicle and JPC-2056-treated groups. Data are presented as mean ± SEM (n = 5 mice per group).

Building on the promising efficacy against *B. duncani* parasites in the lethal infection mouse model, we extended our investigation to assess its effectiveness against *Babesia microti* (ATCC LabS1) in the SCID (severely combined immunodeficient) mouse model of infection following infection with a low dose (1x 10^4^ iRBCs) of parasites. In the group of female mice treated with the vehicle (PEG-400), the highest parasitemia was observed, reaching a maximum of 60% to 70% after 20 days post-infection (DPI-20). Interestingly, daily dosing of JPC-2056 at 30 mg/kg for 10 days (DPI-1 to 10) (**Fig. 3D and E**) successfully eliminated parasites in all infected mice until the end of the study (DPI-45).

### Dihydrotriazines and biguanides, analogs of JPC-2067, exhibit potent broad spectrum antiparasitic activity in vitro

Given the previously reported liabilities of JPC-2067 [43], we sought to evaluate the in vitro activity of dihydrotriazines and biguanides derivatives against several intraerythrocytic parasites for which an in vitro culture system in human erythrocytes has been developed. Due to the diverse culture media used over the years in various laboratories for in vitro culture of intraerythrocytic parasites, the extrapolation of in vitro efficacy and other biological data from one species to another has proven to be a major hurdle in drawing meaningful conclusions across different species. Given the finding that *B. duncani* could be cultured in DFS20 (DMEM-F12 supplemented in 20% FBS), which contains all the nutrients present in RPMI plus three essential ones, putrescine, linoleic acid and lipoic acid, required for *B. duncani*’s growth, we inferred that employing DFS20 for culturing *B. duncani*, *B. divergens*, *B. MO1* and *P. falciparum* strains would facilitate a more precise comparison of the susceptibility of these species and strains under uniform experimental conditions [54, 55]. Consequently, we opted to use the DFS20 culture conditions for conducting a chemical screen aimed at identifying more potent and safer compounds. We screened a total of 79 DHT derivatives (51 dihydrotriazines and 28 biguanides) against three *Babesia* species and *P. falciparum*, which can be continuously propagated in vitro (structures are presented in supplementary data). We assessed inhibition of parasite growth at concentrations of 5, 10, 50 and 100 nM against *B. duncani* (WA-1), *B. divergens* (Rouen87), *B. MO1* (**Fig. 4A-C**) and *P. falciparum* strains (3D7, Dd2 and HB3) (**Fig. 4D-F**). Among the dihydrotriazines and biguanides, biguanides exhibit greater efficacy against *Plasmodium falciparum* compared to *Babesia* species. The compounds were found to be highly effective against the *Plasmodium falciparum* multidrug-resistant strain, Dd2. In an overall fixed concentration screening, dihydrotriazines were found to be more effective against *Plasmodium* species, with approximately 82%, 76%, and 87% of compounds at 100 nM concentration demonstrating effectiveness against *Plasmodium falciparum* 3D7, Dd2, and HB3, respectively (**Fig. 4D-F**). In contrast, the efficacy against *Babesia* species was relatively lower, with effectiveness rates of 67%, 34%, and 60% observed against *Babesia duncani* (WA1), *B. divergens* (Rouen87), and *B. MO1*, respectively. The 9 DHT derivatives with the highest activity against *B. duncani* WA1 also showed broad anti-parasitic activity and were prioritized for further investigations. All 9 compounds showed effective dose-dependent parasite growth inhibition with low nanomolar IC_50_ values ranging from 10 nM to 0.4 nM (**Fig. S5 and Table 2**). The selected compounds were also evaluated for their cytotoxicity profile against human cell lines (HepG2, HCT-116, HEK-293, and HeLa) and displayed low cytotoxicity with therapeutic indices ranging from 4,000 to 100,000 (**Table. 1**). Among these compounds, JPC-2060 emerged as the most broadly effective compound (**Fig. S4**), displaying high potency against *P. falciparum* strains (3D7: IC_50_ ∼1.9 nM; Dd2: IC_50_ ∼0.4 nM; HB3: IC_50_ ∼0.6 nM), *B. duncani* WA1 strain (IC_50_ ∼ 0.8 nM), *B. divergens* Rouen87 strain (IC_50_ ∼3.8 nM); and *B. MO1* (IC_50_ ∼0.5 nM) (**Table 2**).

**Figure 4.**
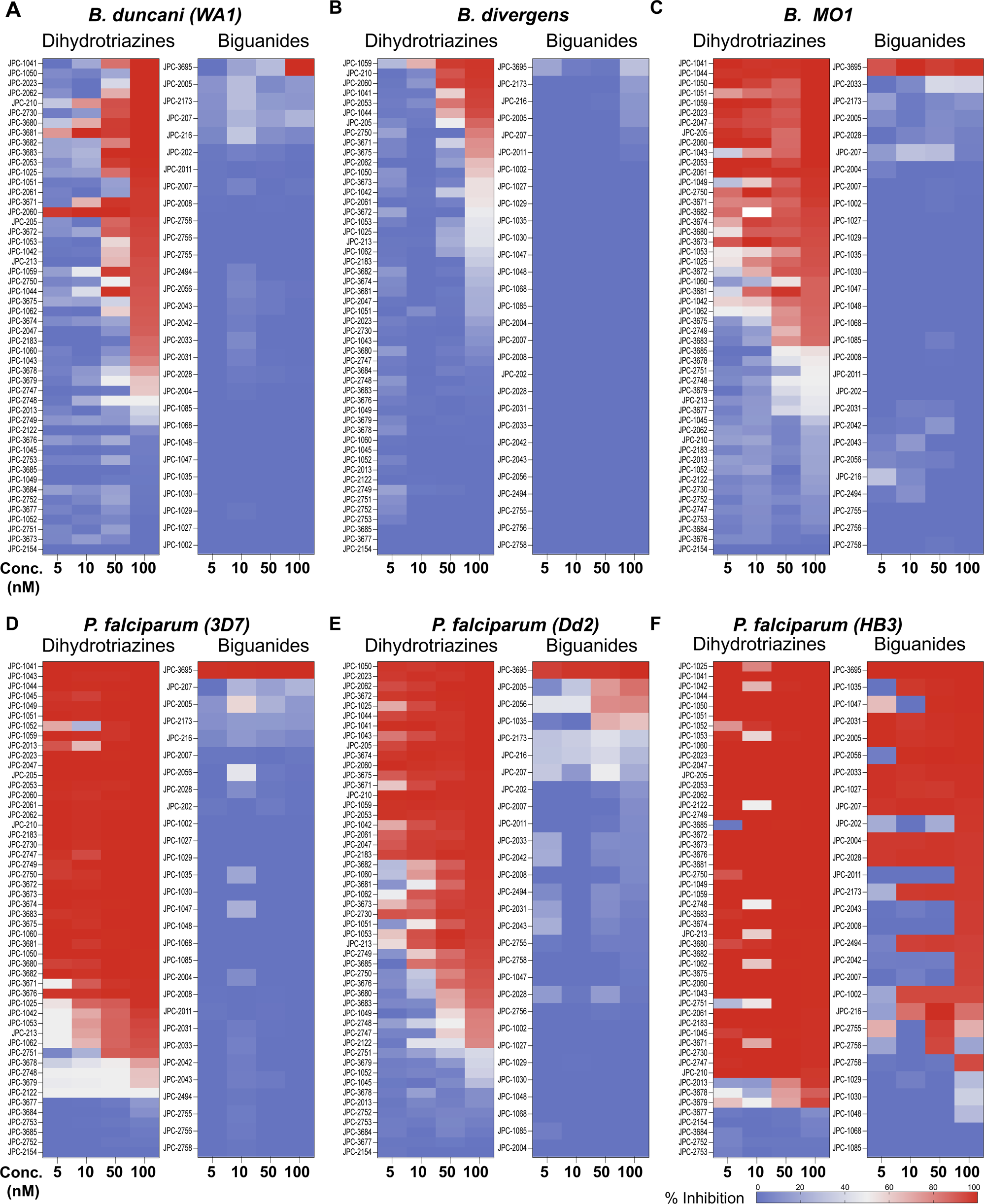
Activity of DHTs and biguanides against *Babesia* and *Plasmodium* species and isolates in vitro. **(A-F)** In vitro efficacy of dihydrotriazines and biguanides against *Babesia* (*B. duncani* WA1**(A)***, B divergens* Rouen87 **(B)**, and *B.MO1* **(C)**) and *Plasmodium* species (3D7 **(D)**, Dd2 **(E)**, and HB3 **(F)**) at 5, 10, 50 and 100 nM drug concentrations. Color-coded heat maps represent mean (n=3) percent inhibition of parasite growth with dark blue representing 100% growth whereas dark red 100% inhibition.

**Table 2.**
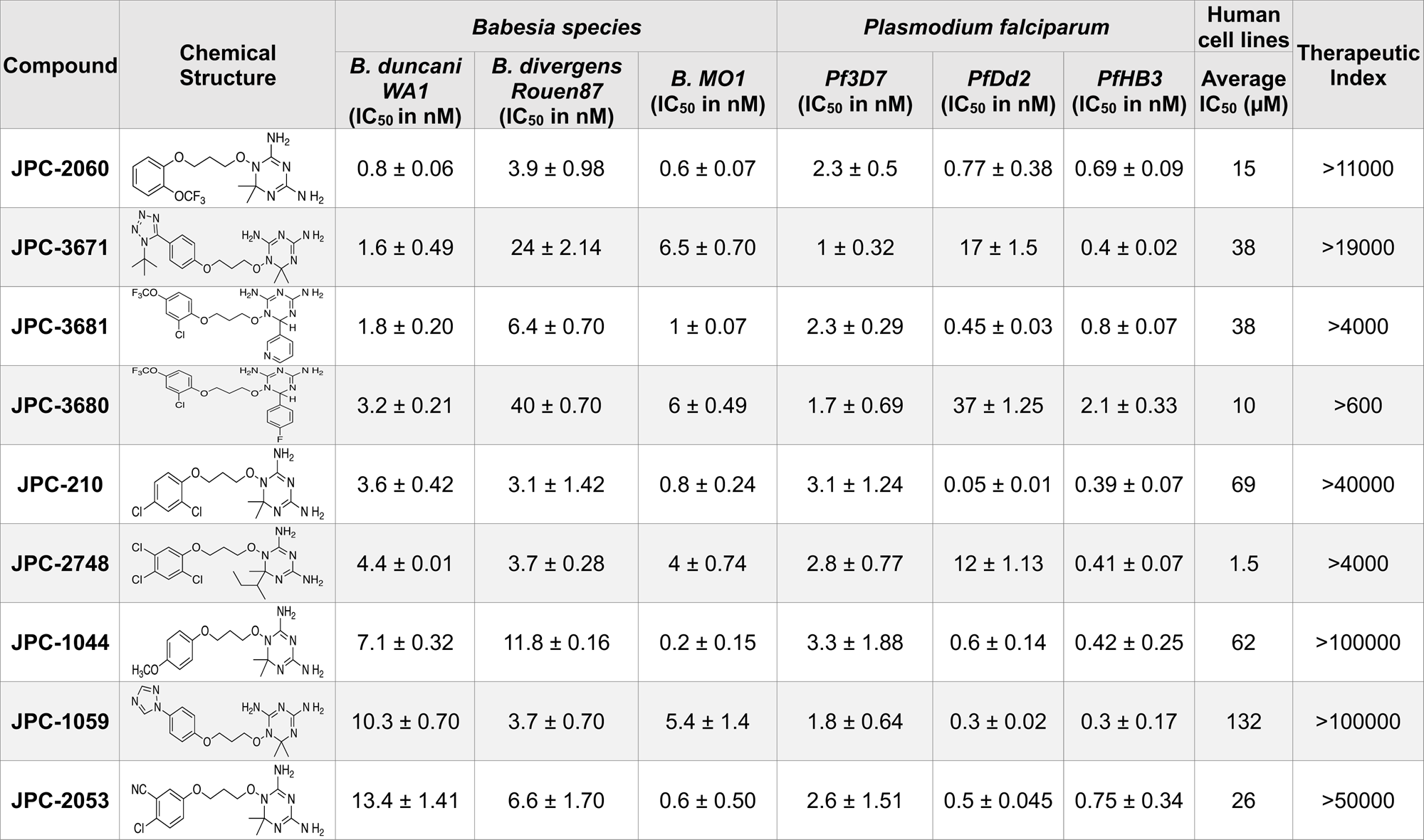
In vitro efficacy and therapeutic indices of dihydrotriazine derivatives.

| Compound | Chemical Structure | *Babesia species* |  |  | *Plasmodium falciparum* |  |  | Human cell lines | Therapeutic Index |
| --- | --- | --- | --- | --- | --- | --- | --- | --- | --- |
|  |  | *B. duncani* WA1<br>(IC<sub>50</sub> in nM) | *B. divergens* Rouen87<br>(IC<sub>50</sub> in nM) | *B. MO1*<br>(IC<sub>50</sub> in nM) | *Pf3D7*<br>(IC<sub>50</sub> in nM) | *PfDd2*<br>(IC<sub>50</sub> in nM) | *PfHB3*<br>(IC<sub>50</sub> in nM) | Average IC<sub>50</sub> (μM) |  |
| JPC-2060 |  | 0.8 ± 0.06 | 3.9 ± 0.98 | 0.6 ± 0.07 | 2.3 ± 0.5 | 0.77 ± 0.38 | 0.69 ± 0.09 | 15 | >11000 |
| JPC-3671 |  | 1.6 ± 0.49 | 24 ± 2.14 | 6.5 ± 0.70 | 1 ± 0.32 | 17 ± 1.5 | 0.4 ± 0.02 | 38 | >19000 |
| JPC-3681 |  | 1.8 ± 0.20 | 6.4 ± 0.70 | 1 ± 0.07 | 2.3 ± 0.29 | 0.45 ± 0.03 | 0.8 ± 0.07 | 38 | >4000 |
| JPC-3680 |  | 3.2 ± 0.21 | 40 ± 0.70 | 6 ± 0.49 | 1.7 ± 0.69 | 37 ± 1.25 | 2.1 ± 0.33 | 10 | >600 |
| JPC-210 |  | 3.6 ± 0.42 | 3.1 ± 1.42 | 0.8 ± 0.24 | 3.1 ± 1.24 | 0.05 ± 0.01 | 0.39 ± 0.07 | 69 | >40000 |
| JPC-2748 |  | 4.4 ± 0.01 | 3.7 ± 0.28 | 4 ± 0.74 | 2.8 ± 0.77 | 12 ± 1.13 | 0.41 ± 0.07 | 1.5 | >4000 |
| JPC-1044 |  | 7.1 ± 0.32 | 11.8 ± 0.16 | 0.2 ± 0.15 | 3.3 ± 1.88 | 0.6 ± 0.14 | 0.42 ± 0.25 | 62 | >100000 |
| JPC-1059 |  | 10.3 ± 0.70 | 3.7 ± 0.70 | 5.4 ± 1.4 | 1.8 ± 0.64 | 0.3 ± 0.02 | 0.3 ± 0.17 | 132 | >100000 |
| JPC-2053 |  | 13.4 ± 1.41 | 6.6 ± 1.70 | 0.6 ± 0.50 | 2.6 ± 1.51 | 0.5 ± 0.045 | 0.75 ± 0.34 | 26 | >50000 |

## Discussion

The DHFR-TS enzymes of *Babesia* species exhibit notable differences compared to those of *P. falciparum*, particularly in size and amino acid composition. Unlike the longer *P. falciparum* enzyme, *Babesia* DHFR-TS enzymes are comparatively smaller due to the absence of large insertions in their sequences. However, they share high similarity with the *P. falciparum* enzyme, suggesting a close evolutionary relationship within the *Apicomplexa* phylum.

The in vitro assessment of the antifolate JPC-2067 and its pro-drug JPC-2056 revealed potent inhibitory activity against *B. duncani*, with nanomolar IC_50_ values. Importantly, these compounds exhibited high selectivity with minimal cytotoxicity against human cell lines. Furthermore, JPC-2067 effectively inhibited *Babesia* DHFR-TS enzymes, underscoring its mechanism of action. In vivo studies using JPC-2056 demonstrated efficacy against *B. duncani* in a murine model of babesiosis. Although complete parasite clearance was not achieved, treatment with JPC-2056 resulted in a significant delay in parasite emergence and reduced peak parasitemia. Moreover, JPC-2056 showed remarkable efficacy against *B. microti* in a SCID mouse model of babesiosis, leading to the elimination of parasites without observable adverse effects.

Expanding on the efficacy of JPC-2067, we sought to evaluate structurally related dihydrotriazines and biguanides for their antibabesial as well as antimalarial activity. However, comparing the efficacy of various compounds across different species and strains poses challenges in drug susceptibility testing. Variations in culture conditions can introduce confounding factors, making it difficult to draw accurate conclusions. Our present study addressed this challenge by employing DFS20, a culture medium that provides uniform nutrient supplementation, including essential elements such as putrescine, linoleic acid, and lipoic acid crucial for the growth of diverse parasites. By utilizing DFS20 as a standardized culture medium for culturing *B. duncani*, *B. divergens*, *B. MO1*, and *P. falciparum* strains, we have created a platform conducive to more reliable cross-species and cross-strain comparisons of drug susceptibility. Using these standardized conditions, we screened a new library of 51 dihydrotriazines and 28 biguanides against all these species. Nine promising candidates were identified with IC_50_ values below 10 nM against all species and strains tested, and no toxicity at concentrations up to 900 µM. Both multi-drug resistant and drug-sensitive *P. falciparum* strains were sensitive to the tested compounds. Among these candidates, JPC-2060 emerged as the most promising candidate for further investigation. However, treating mice infected with *B. duncani* with prodrugs for six of these candidates and all biguanides at 30 mg/kg for five days did not confer protection against this infection. Further optimization, including structural modifications and synthesis of more bioactive analogs, is thus necessary to improve bioavailability and extend efficacy studies to live animal models.

Our screening of analogs against *B. duncani* revealed a notable structure-activity relationship, particularly with the 2-Chloro, 4-Trifluoromethoxy analogs. By altering the substitution on the dihydrotriazine ring from dimethyl to aryl, we found that adding an electron deficient aryl group at C-6 increased activity in the case of 6-(3-pyridyl) JPC-3681 and 6-(4-fluorophenyl) JPC-3680. Substituting the aryl group with an alkyl group, like 6-propyl JPC-3682 or a different pyridine, like 6-(2-pyridyl) JPC-3683, maintained some activity. Conversely, compounds with electron-rich or neutral aryl groups, such as 6-(4-methoxyphenyl) JPC-3678 and 6-phenyl JPC-3679 exhibited decreased activity. Additionally, appending nitrogen heterocycles on the terminal aryl group proved beneficial, as observed with 4-(1-(t-butyl)-1H Tetrazol-5-yl) JPC-3671 or 4-(1H-1,2,4-triazol-1-yl) JPC-1059. These findings suggest avenues for further exploration by either combining or delving deeper into these structural regions.

In conclusion, our study underscores the importance of standardized culture conditions in facilitating accurate comparisons of drug susceptibility across different parasitic species and strains. The identification of highly potent pan-antiparasitic drugs targeting DHFR-TS enzymes represents a significant advancement in the quest for effective treatments against malaria and babesiosis. Future research should focus on further optimization of these early lead compounds to identify candidates with biological activity in animal models of babesiosis and malaria.

## Materials and Methods

Unless otherwise stated, chemicals were purchased from commercial suppliers and used as received. WR99210, JPC-derivatives, and DHT derivatives with purity ≥ 95%, as measured by reversed-phase high-performance liquid chromatography (HPLC), were procured from Jacobus Pharmaceuticals.

### Animal studies

Immunocompetent C3H/HeJ mice were procured from The Jackson Laboratory, and immunocompromised CB17/SCID mice were obtained from Envigo. All animal experiments strictly adhered to Yale University’s institutional guidelines for the care and use of laboratory animals and were conducted under the approval of a protocol sanctioned by the Institutional Animal Care and Use Committees (IACUC) at Yale University.

### Chemical synthesis of dihydrotriazines and biguanides

The dihydrotriazines were prepared as previously described by Jensen *et al*. [56]. The majority of the dihydrotriazines were made by acidic condensation of the related pro-drug biguanide with the corresponding ketone, with acetone (Y and Z = CH3) as the standard as shown in step (g) as described in scheme (**Fig. S5**). The biguanides were prepared as described both in the Jensen paper and as more fully described in the patent literature [57, 58]. Compound identity and purity were confirmed by LC/MS. The purity of all compounds at the time of testing was greater than 95%. The structures of the compounds assayed, the observed molecular ions, and the Chemical Abstract Service numbers (CAS #) for previously reported compounds can be found in Table S1 and S2.

### In-vitro parasite culture of Babesia duncani and Plasmodium falciparum

Continuous propagation of *B. duncani* in human RBCs in vitro were carried out as first reported earlier by Abraham *et al.* and further optimized [20, 49]. Briefly, parasites cultured in a complete HL-1 medium (Base medium of HL-1 (Lonza, 344017) supplemented with 20% heat-inactivated FBS, 2% 50X HT Media Supplement Hybrid-MaxTM (Sigma, H0137), 1% 200 mM L-Glutamine (Gibco, 25030-081), 1% 100X Penicillin/Streptomycin (Gibco, 15240-062) and 1% 10 mg/mL Gentamicin (Gibco, 15710-072)) in 5% hematocrit A+ RBCs and maintained at 37°C in a humidified chamber containing 2% O_2_, 5% CO_2_, and 93% N_2_. *P. falciparum* (3D7, Dd2, and HB3) parasites were cultured in 3-5% human O^+^ RBCs in RPMI-1640 (Gibco, 110875093) complete medium containing 0.5% Albumax-II (Invitrogen), 2 mM L-glutamine, 50 mg/L hypoxanthine, 25 mM HEPES, 0.225% NaHCO_3_ and 10 mg/mL gentamycin. Cultures were maintained at 37°C and gassed with a sterile mixture of 4% O_2_, 5% CO_2,_ and 91% N2. The culture medium was changed daily, and parasitemia was monitored by light microscope examination of Giemsa-stained thin-blood smears.

### Assessment of drug cytotoxicity on human cell lines

HeLa, HepG2, HEK, and HCT116 cell lines were obtained from the American Type Culture Collection (ATCC) and maintained in Dulbecco’s modified Eagle’s medium (DMEM) (Invitrogen 11995-065) containing 25 mM glucose, 1 mM sodium pyruvate and supplemented with 5 mM HEPES, 10% FBS and penicillin-streptomycin (50 U/mL penicillin, 50 µg/mL streptomycin). Cells were seeded in a 96-well tissue culture plate (20,000 cells per well), allowed to adhere for 24 hours, after which they were treated with a 12-step 2-fold serial dilution of each drug starting at 1 mM as the highest final concentration. Cells maintained in medium supplemented with either 0.1% or 10% DMSO were used as negative and positive vehicle controls, respectively. The plates were incubated at 37°C for 72 hours, after which each well was incubated with 0.5 mg/mL of MTT reagent (MP Chemicals #02102227) for 4 hours in the dark at 37°C. The formazan crystals formed by living cells were solubilized with the addition of 100 µl of DMSO to each well. We measured the OD_590nm_ using a SpectraMax plate reader. From the obtained OD values, percent cell viability was estimated by normalizing to the mean of 10% DMSO wells (set as 100% toxicity) and mean of the vehicle control wells (set as 0% toxicity). Dose-response curves were plotted using GraphPad Prism version 9.1.2.

### Enzyme inhibition assays

The inhibition of DHFR activity with selected JPC derivatives was measured as previously described [52]. Briefly, the purified enzyme (0.1 μM) **(Fig. S3**) was incubated with increasing concentrations of JPC-2067 in a reaction buffer containing 300 μM DHF and 300 μM NADPH. OD_340_ reduction rate (OD_340_/min) was measured for further calculation. The % inhibition was calculated using the formula:

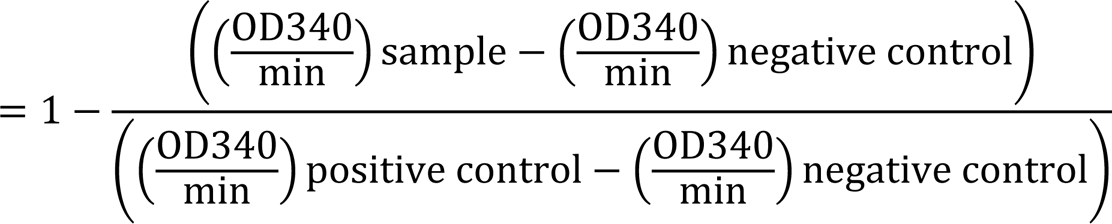

The IC_50_ was determined from a sigmoidal dose-response curve using GraphPad Prism version 9.1.2.

### In vitro drug efficacy

The effect of DHT derivatives on the intraerythrocytic development of *P. falciparum* and *B. duncani* and determination of IC_50_ values were conducted as previously reported [49, 59]. Briefly, the parasites cultured in vitro (0.1% parasitemia with 5% hematocrit in complete DMEM-F12 medium or RPMI-1640) were treated with various concentrations of the compounds in a 96-well plate and incubated at 37°C for either 60 h in the case of *B. duncani* or 72 h in the case of *P. falciparum.* Parasitemia was then determined using the SYBR Green-I method by adding an equal volume of parasite culture to the lysis buffer (0.008% saponin, 0.08% Triton-X-100, 20 mM Tris-HCl (pH = 7.5) and 5 mM EDTA) containing SYBR Green-I (0.01%). The mixture was then incubated at 37°C for 1 h in the dark. The fluorescence was measured at 480 nm (excitation) and 540 nm (emission) using a BioTek Synergy™ Mx Microplate Reader. Background fluorescence from uninfected erythrocytes maintained in complete HL-1and RPMI-1640 medium was subtracted from each sample and the 50% inhibitory concentration (IC_50_) of the drug was determined by plotting drug concentrations versus parasite growth using GraphPad Prism 9.2.1. Data are shown as mean ± SD from two independent experiments each with biological triplicates.

### In vivo drug efficacy

Both C3H/HeJ and CB17/SCID mice aged 5-6 weeks (4 to 5 mice per group in each of the studies described herein) were injected intravenously with low dose (10^4^ iRBCs) of *B duncani* (WA-1) and *B. microti* (LabS1) parasites respectively. The infected mice were treated by oral gavage over the 10 days (DPI 1 to 10). During the treatment, each mouse received 100 µL of either vehicle (PEG-400), JPC-2056 (10 mg/kg) or JPC-2056 (30 mg/kg). Percent parasitemia was determined by counting at least 3000 RBCs from Giemsa-stained thick blood smears.

#### Statistical analysis

Datasets were analyzed with GraphPad Prism version 9.1.2. Statistical significances were determined using unpaired t-tests.

---

**Figure S1.**
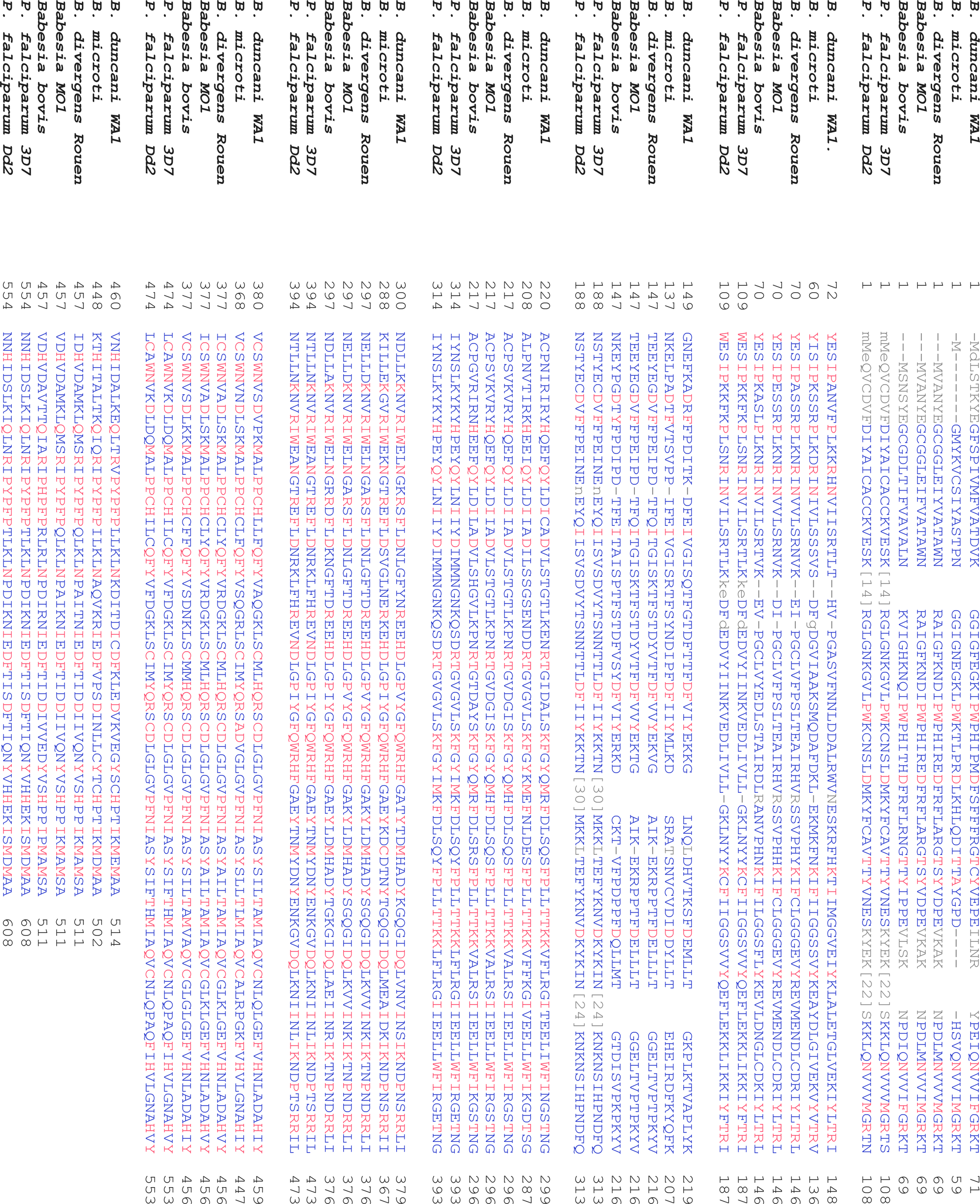
Alignment of the full-length sequences of DHFR-TS from *Babesia* and *Plasmodium* parasites.

**Figure S2.**
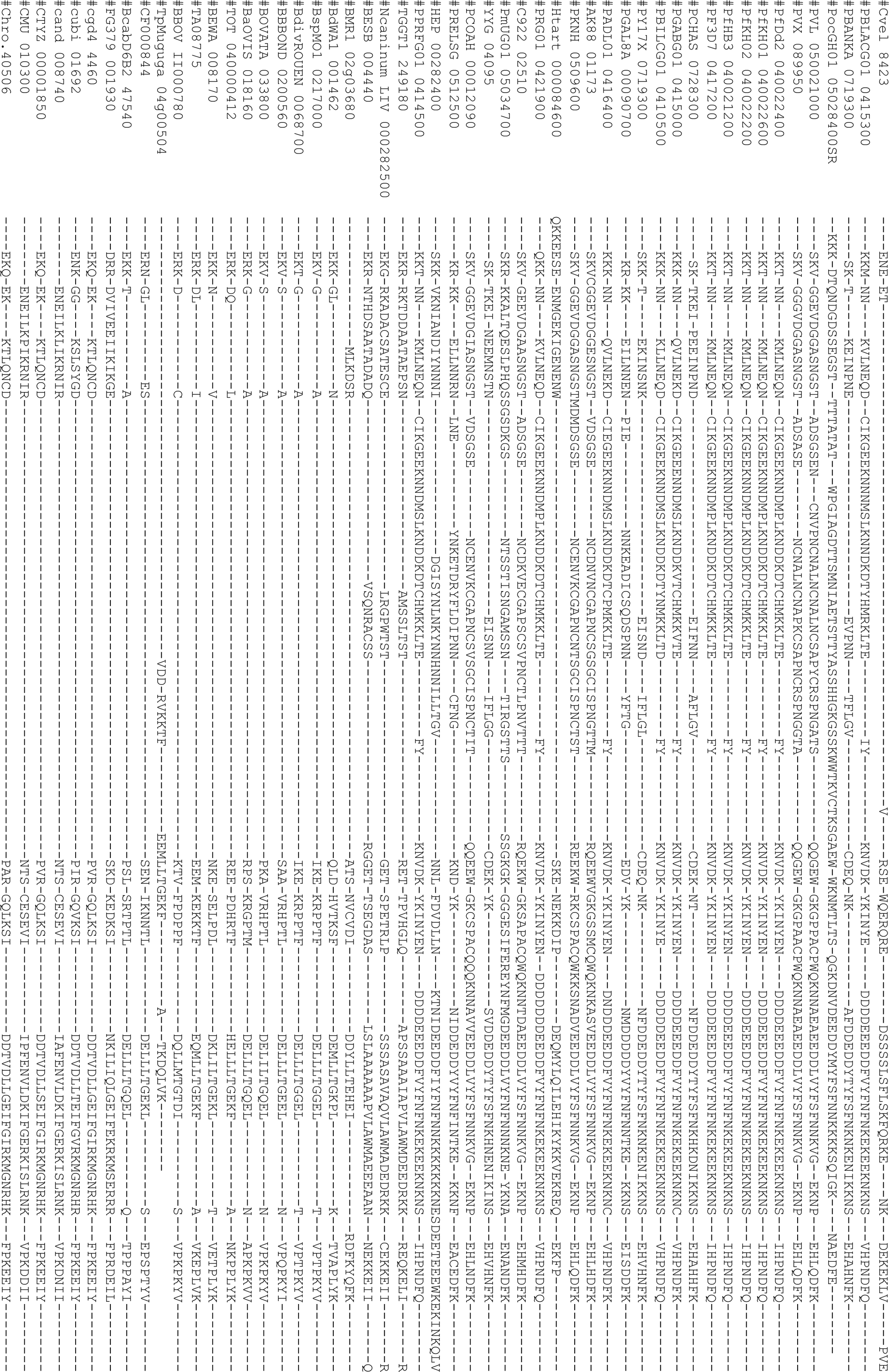
Alignment of the sequence linking the DHFR and TS domains in the DHFR-TS bifunctional enzymes from 52 apicomplexan parasites (ID, species name, protein length are provided in Suppl Data).

**Figure S3.**
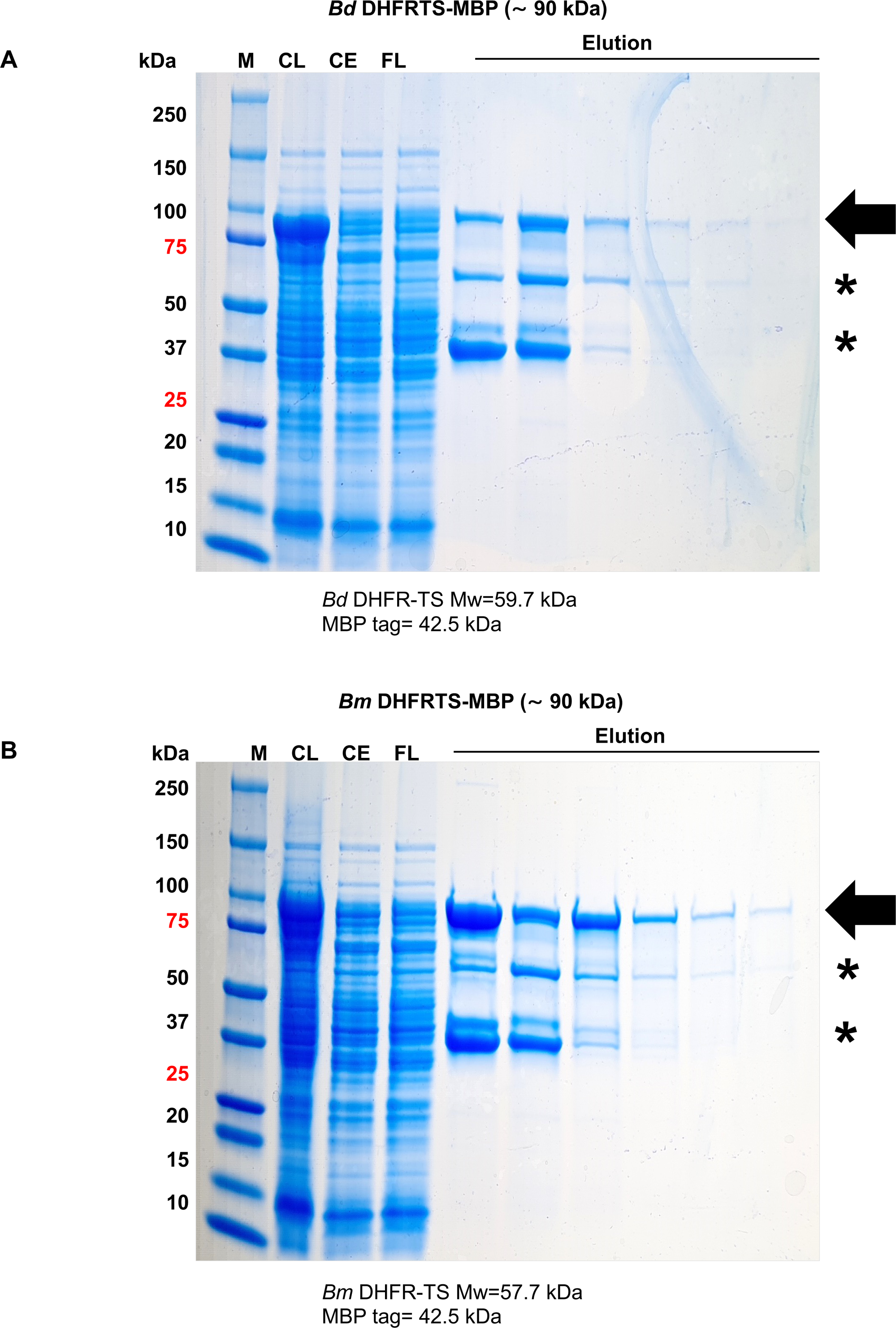
Coomassie blue-stained SDS-PAGE gels showing recombinant *B. duncani* **(A)** and *B. microti* -DHFR-TS **(B)** enzymes purified from E. coli cell extracts using an amylose resin. Total cell extracts (CE), flow-through (FT), and elution fractions along with protein standard marker (M) were loaded on a gradient SDS-PAGE (4-10%). The purified Bd and Bm-DHFR-TS fused with the MBP-tag migrate at approximately ∼90 kDa size. Purified enzymes were eluted with elution buffer containing 10 mM maltose. Amylose elution fractions were loaded onto MTX-agarose resin and purified DHFR-TS enzymes were eluted using 2 mM dihydrofolic acid (DHF). Maltose or DHF residuals were removed by dialysis or using a PD-10 column (GE Healthcare).

**Figure S4.**
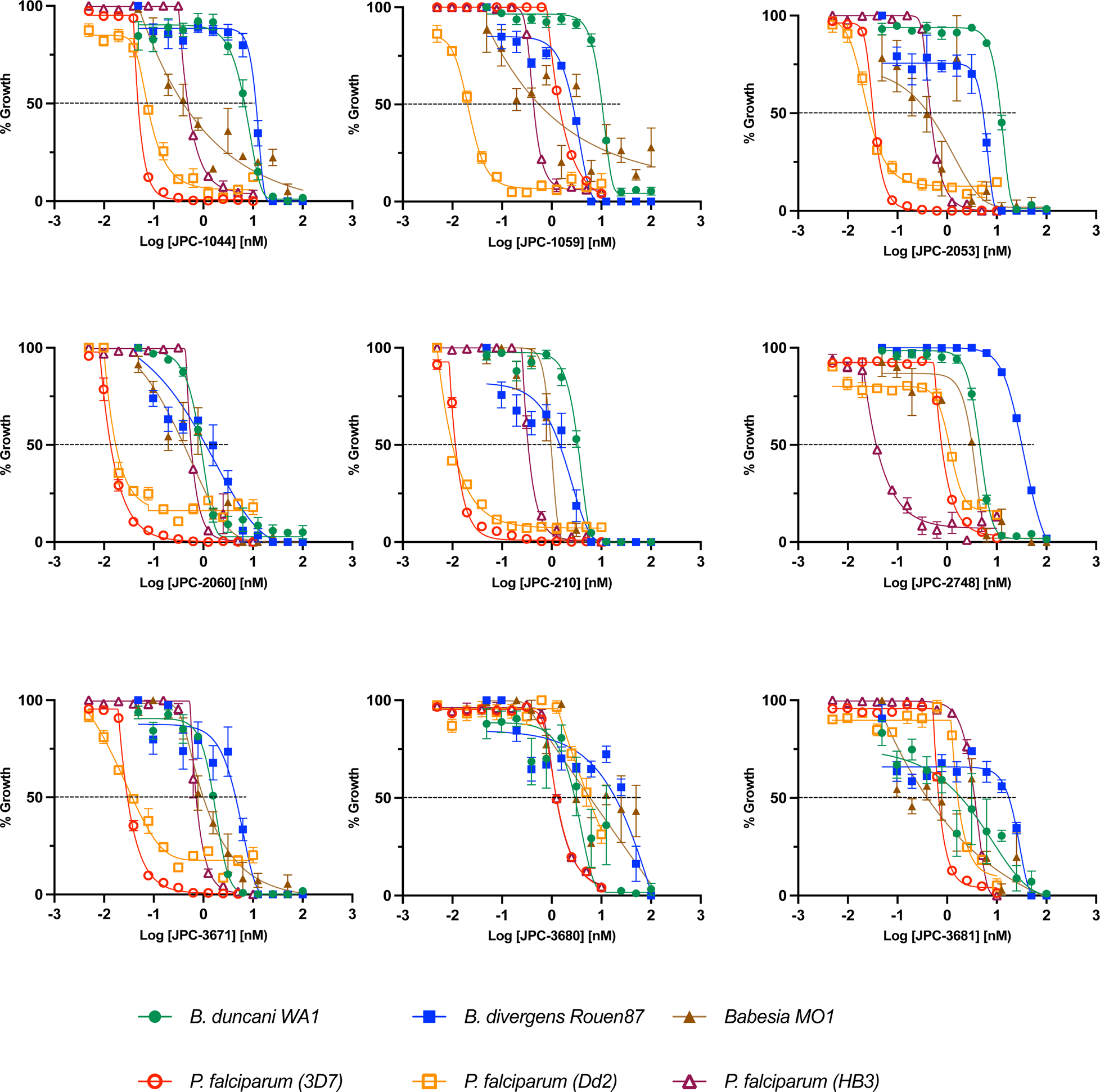
Efficacy of shortlisted DHT derivatives. Dose-response sigmoidal semilogarithmic graphs with parasite growth inhibition (y-axis) versus the log concentration of 9 DHT derivatives (x-axis) against *Babesia* (*B. duncani WA1*(green)*, Bdiv Rouen87* (blue)*, and B. spMO1*(brown)) and *Plasmodium* species (3D7 (red), Dd2 (yellow), and HB3(purple)).

**Figure S5.**
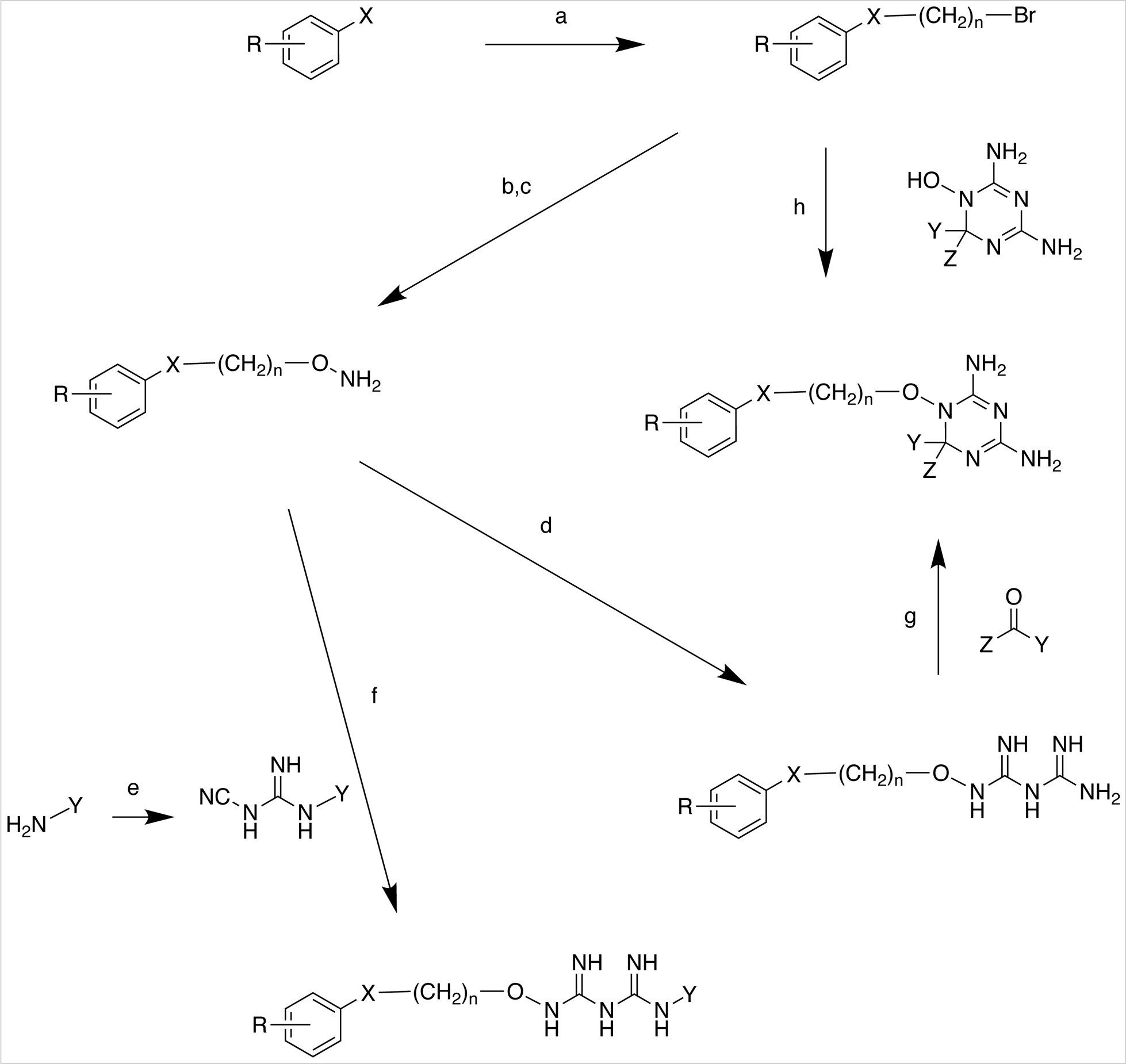
Scheme for the chemical synthesis of dihydrotriazines and biguanides. Reagents: (a) 1,3-dibromopropane (for n = 3), NaOH, tetrabutylammonium hydrogen sulfate; (b) AcNHOH, NaOH, or KOH; alcoholic solvent; (c) concentrated HCl, MeOH; (d) dicyandiamide, aqueous EtOH, heat, and then aqueous NaOH to neutralize; (e) sodium dicyanamide, HCl, alcoholic solvent, heat; (f) EtOAc, heat; (g) HCl, MeOH; (h) room temp, DMF.

**Table S1.**
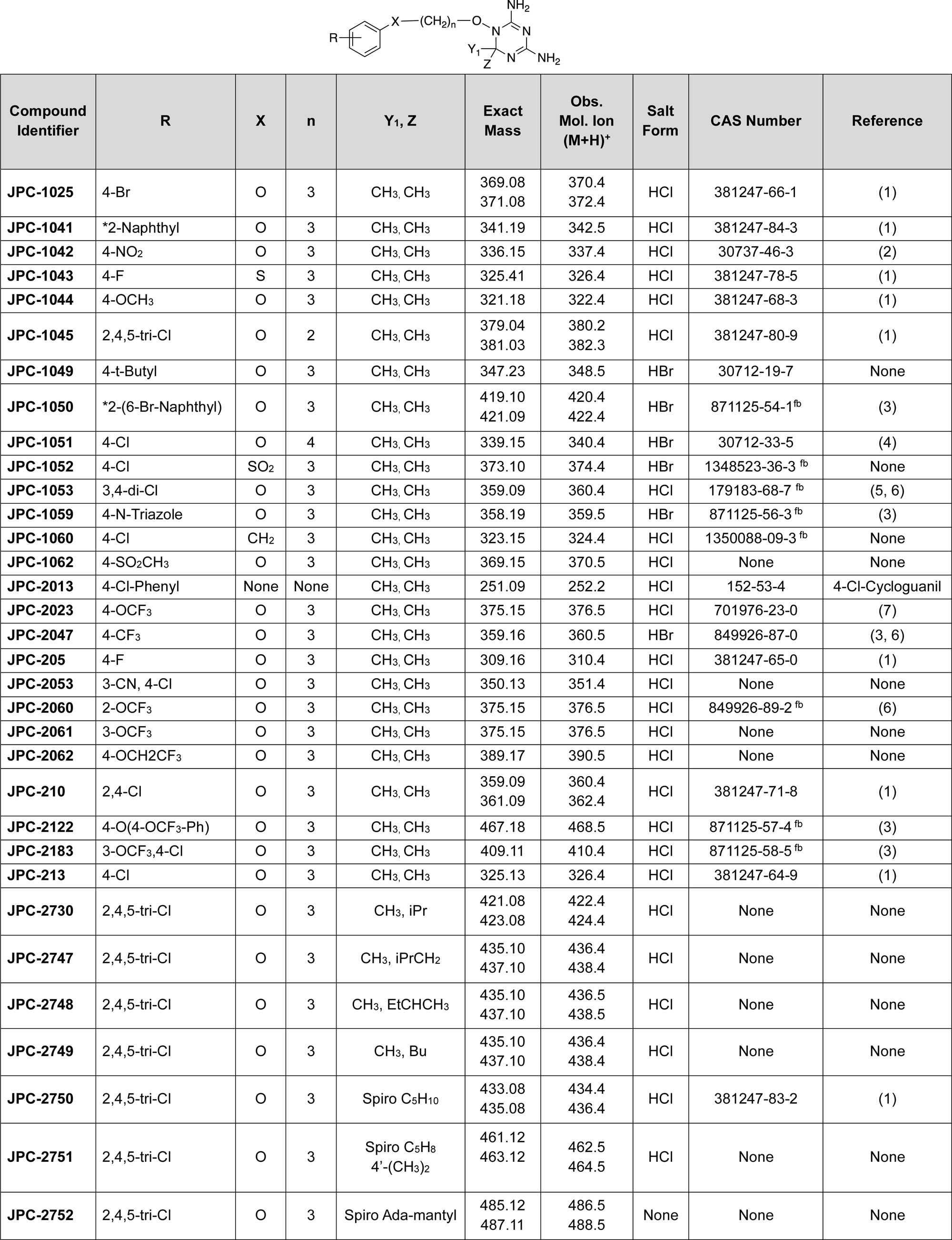
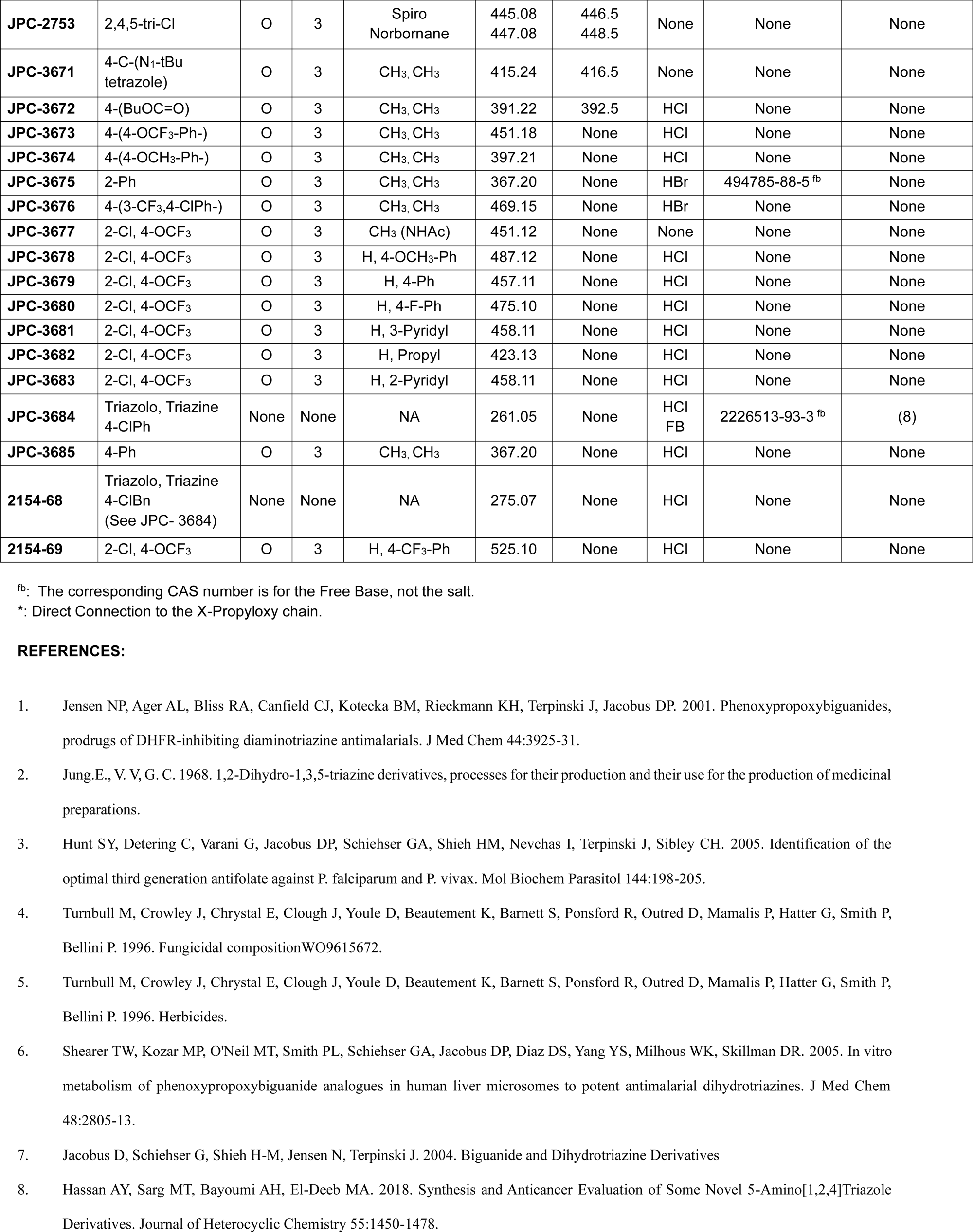
List of Dihydrotriazine derivatives with Chemical Abstract System (CAS) number, functional groups, and corresponding references.

| Compound Identifier | R | X | n | Y<sub>1</sub>, Z | Exact Mass | Obs. Mol. Ion (M+H)<sup>+</sup> | Salt Form | CAS Number | Reference |
| --- | --- | --- | --- | --- | --- | --- | --- | --- | --- |
| JPC-1025 | 4-Br | O | 3 | CH<sub>3</sub>, CH<sub>3</sub> | 369.08<br>371.08 | 370.4<br>372.4 | HCl | 381247-66-1 | (1) |
| JPC-1041 | *2-Naphthyl | O | 3 | CH<sub>3</sub>, CH<sub>3</sub> | 341.19 | 342.5 | HCl | 381247-84-3 | (1) |
| JPC-1042 | 4-NO<sub>2</sub> | O | 3 | CH<sub>3</sub>, CH<sub>3</sub> | 336.15 | 337.4 | HCl | 30737-46-3 | (2) |
| JPC-1043 | 4-F | S | 3 | CH<sub>3</sub>, CH<sub>3</sub> | 325.41 | 326.4 | HCl | 381247-78-5 | (1) |
| JPC-1044 | 4-OCH<sub>3</sub> | O | 3 | CH<sub>3</sub>, CH<sub>3</sub> | 321.18 | 322.4 | HCl | 381247-68-3 | (1) |
| JPC-1045 | 2,4,5-tri-Cl | O | 2 | CH<sub>3</sub>, CH<sub>3</sub> | 379.04<br>381.03 | 380.2<br>382.3 | HCl | 381247-80-9 | (1) |
| JPC-1049 | 4-t-Butyl | O | 3 | CH<sub>3</sub>, CH<sub>3</sub> | 347.23 | 348.5 | HBr | 30712-19-7 | None |
| JPC-1050 | *2-(6-Br-Naphthyl) | O | 3 | CH<sub>3</sub>, CH<sub>3</sub> | 419.10<br>421.09 | 420.4<br>422.4 | HBr | 871125-54-1<sup>fb</sup> | (3) |
| JPC-1051 | 4-Cl | O | 4 | CH<sub>3</sub>, CH<sub>3</sub> | 339.15 | 340.4 | HBr | 30712-33-5 | (4) |
| JPC-1052 | 4-Cl | SO<sub>2</sub> | 3 | CH<sub>3</sub>, CH<sub>3</sub> | 373.10 | 374.4 | HBr | 1348523-36-3<sup>fb</sup> | None |
| JPC-1053 | 3,4-di-Cl | O | 3 | CH<sub>3</sub>, CH<sub>3</sub> | 359.09 | 360.4 | HCl | 179183-68-7<sup>fb</sup> | (5, 6) |
| JPC-1059 | 4-N-Triazole | O | 3 | CH<sub>3</sub>, CH<sub>3</sub> | 358.19 | 359.5 | HBr | 871125-56-3<sup>fb</sup> | (3) |
| JPC-1060 | 4-Cl | CH<sub>2</sub> | 3 | CH<sub>3</sub>, CH<sub>3</sub> | 323.15 | 324.4 | HCl | 1350088-09-3<sup>fb</sup> | None |
| JPC-1062 | 4-SO<sub>2</sub>CH<sub>3</sub> | O | 3 | CH<sub>3</sub>, CH<sub>3</sub> | 369.15 | 370.5 | HCl | None | None |
| JPC-2013 | 4-Cl-Phenyl | None | None | CH<sub>3</sub>, CH<sub>3</sub> | 251.09 | 252.2 | HCl | 152-53-4 | 4-Cl-Cycloguanil |
| JPC-2023 | 4-OCF<sub>3</sub> | O | 3 | CH<sub>3</sub>, CH<sub>3</sub> | 375.15 | 376.5 | HCl | 701976-23-0 | (7) |
| JPC-2047 | 4-CF<sub>3</sub> | O | 3 | CH<sub>3</sub>, CH<sub>3</sub> | 359.16 | 360.5 | HBr | 849926-87-0 | (3, 6) |
| JPC-205 | 4-F | O | 3 | CH<sub>3</sub>, CH<sub>3</sub> | 309.16 | 310.4 | HCl | 381247-65-0 | (1) |
| JPC-2053 | 3-CN, 4-Cl | O | 3 | CH<sub>3</sub>, CH<sub>3</sub> | 350.13 | 351.4 | HCl | None | None |
| JPC-2060 | 2-OCF<sub>3</sub> | O | 3 | CH<sub>3</sub>, CH<sub>3</sub> | 375.15 | 376.5 | HCl | 849926-89-2<sup>fb</sup> | (6) |
| JPC-2061 | 3-OCF<sub>3</sub> | O | 3 | CH<sub>3</sub>, CH<sub>3</sub> | 375.15 | 376.5 | HCl | None | None |
| JPC-2062 | 4-OCH<sub>2</sub>CF<sub>3</sub> | O | 3 | CH<sub>3</sub>, CH<sub>3</sub> | 389.17 | 390.5 | HCl | None | None |
| JPC-210 | 2,4-Cl | O | 3 | CH<sub>3</sub>, CH<sub>3</sub> | 359.09<br>361.09 | 360.4<br>362.4 | HCl | 381247-71-8 | (1) |
| JPC-2122 | 4-O(4-OCF<sub>3</sub>-Ph) | O | 3 | CH<sub>3</sub>, CH<sub>3</sub> | 467.18 | 468.5 | HCl | 871125-57-4<sup>fb</sup> | (3) |
| JPC-2183 | 3-OCF<sub>3</sub>,4-Cl | O | 3 | CH<sub>3</sub>, CH<sub>3</sub> | 409.11 | 410.4 | HCl | 871125-58-5<sup>fb</sup> | (3) |
| JPC-213 | 4-Cl | O | 3 | CH<sub>3</sub>, CH<sub>3</sub> | 325.13 | 326.4 | HCl | 381247-64-9 | (1) |
| JPC-2730 | 2,4,5-tri-Cl | O | 3 | CH<sub>3</sub>, iPr | 421.08<br>423.08 | 422.4<br>424.4 | HCl | None | None |
| JPC-2747 | 2,4,5-tri-Cl | O | 3 | CH<sub>3</sub>, iPrCH<sub>2</sub> | 435.10<br>437.10 | 436.4<br>438.4 | HCl | None | None |
| JPC-2748 | 2,4,5-tri-Cl | O | 3 | CH<sub>3</sub>, EtCHCH<sub>3</sub> | 435.10<br>437.10 | 436.5<br>438.5 | HCl | None | None |
| JPC-2749 | 2,4,5-tri-Cl | O | 3 | CH<sub>3</sub>, Bu | 435.10<br>437.10 | 436.4<br>438.4 | HCl | None | None |
| JPC-2750 | 2,4,5-tri-Cl | O | 3 | Spiro C<sub>5</sub>H<sub>10</sub> | 433.08<br>435.08 | 434.4<br>436.4 | HCl | 381247-83-2 | (1) |
| JPC-2751 | 2,4,5-tri-Cl | O | 3 | Spiro C<sub>5</sub>H<sub>8</sub><br>4'-(CH<sub>3</sub>)<sub>2</sub> | 461.12<br>463.12 | 462.5<br>464.5 | HCl | None | None |
| JPC-2752 | 2,4,5-tri-Cl | O | 3 | Spiro Ada-mantyl | 485.12<br>487.11 | 486.5<br>488.5 | None | None | None |
| **JPC-2753** | 2,4,5-tri-Cl | O | 3 | Spiro<br>Norbornane | 445.08<br>447.08 | 446.5<br>448.5 | None | None | None |
| **JPC-3671** | 4-C-(N<sub>1</sub>-tBu<br>tetrazole) | O | 3 | CH<sub>3</sub>, CH<sub>3</sub> | 415.24 | 416.5 | None | None | None |
| **JPC-3672** | 4-(BuOC=O) | O | 3 | CH<sub>3</sub>, CH<sub>3</sub> | 391.22 | 392.5 | HCl | None | None |
| **JPC-3673** | 4-(4-OCF<sub>3</sub>-Ph-) | O | 3 | CH<sub>3</sub>, CH<sub>3</sub> | 451.18 | None | HCl | None | None |
| **JPC-3674** | 4-(4-OCH<sub>3</sub>-Ph-) | O | 3 | CH<sub>3</sub>, CH<sub>3</sub> | 397.21 | None | HCl | None | None |
| **JPC-3675** | 2-Ph | O | 3 | CH<sub>3</sub>, CH<sub>3</sub> | 367.20 | None | HBr | 494785-88-5<sup>fb</sup> | None |
| **JPC-3676** | 4-(3-CF<sub>3</sub>,4-ClPh-) | O | 3 | CH<sub>3</sub>, CH<sub>3</sub> | 469.15 | None | HBr | None | None |
| **JPC-3677** | 2-Cl, 4-OCF<sub>3</sub> | O | 3 | CH<sub>3</sub> (NHAc) | 451.12 | None | None | None | None |
| **JPC-3678** | 2-Cl, 4-OCF<sub>3</sub> | O | 3 | H, 4-OCH<sub>3</sub>-Ph | 487.12 | None | HCl | None | None |
| **JPC-3679** | 2-Cl, 4-OCF<sub>3</sub> | O | 3 | H, 4-Ph | 457.11 | None | HCl | None | None |
| **JPC-3680** | 2-Cl, 4-OCF<sub>3</sub> | O | 3 | H, 4-F-Ph | 475.10 | None | HCl | None | None |
| **JPC-3681** | 2-Cl, 4-OCF<sub>3</sub> | O | 3 | H, 3-Pyridyl | 458.11 | None | HCl | None | None |
| **JPC-3682** | 2-Cl, 4-OCF<sub>3</sub> | O | 3 | H, Propyl | 423.13 | None | HCl | None | None |
| **JPC-3683** | 2-Cl, 4-OCF<sub>3</sub> | O | 3 | H, 2-Pyridyl | 458.11 | None | HCl | None | None |
| **JPC-3684** | Triazolo, Triazine<br>4-ClPh | None | None | NA | 261.05 | None | HCl<br>FB | 2226513-93-3<sup>fb</sup> | (8) |
| **JPC-3685** | 4-Ph | O | 3 | CH<sub>3</sub>, CH<sub>3</sub> | 367.20 | None | HCl | None | None |
| **2154-68** | Triazolo, Triazine<br>4-ClBn<br>(See JPC- 3684) | None | None | NA | 275.07 | None | HCl | None | None |
| **2154-69** | 2-Cl, 4-OCF<sub>3</sub> | O | 3 | H, 4-CF<sub>3</sub>-Ph | 525.10 | None | HCl | None | None |

<sup>fb</sup>: The corresponding CAS number is for the Free Base, not the salt.
\*: Direct Connection to the X-Propyloxy chain.

**Table S2.**
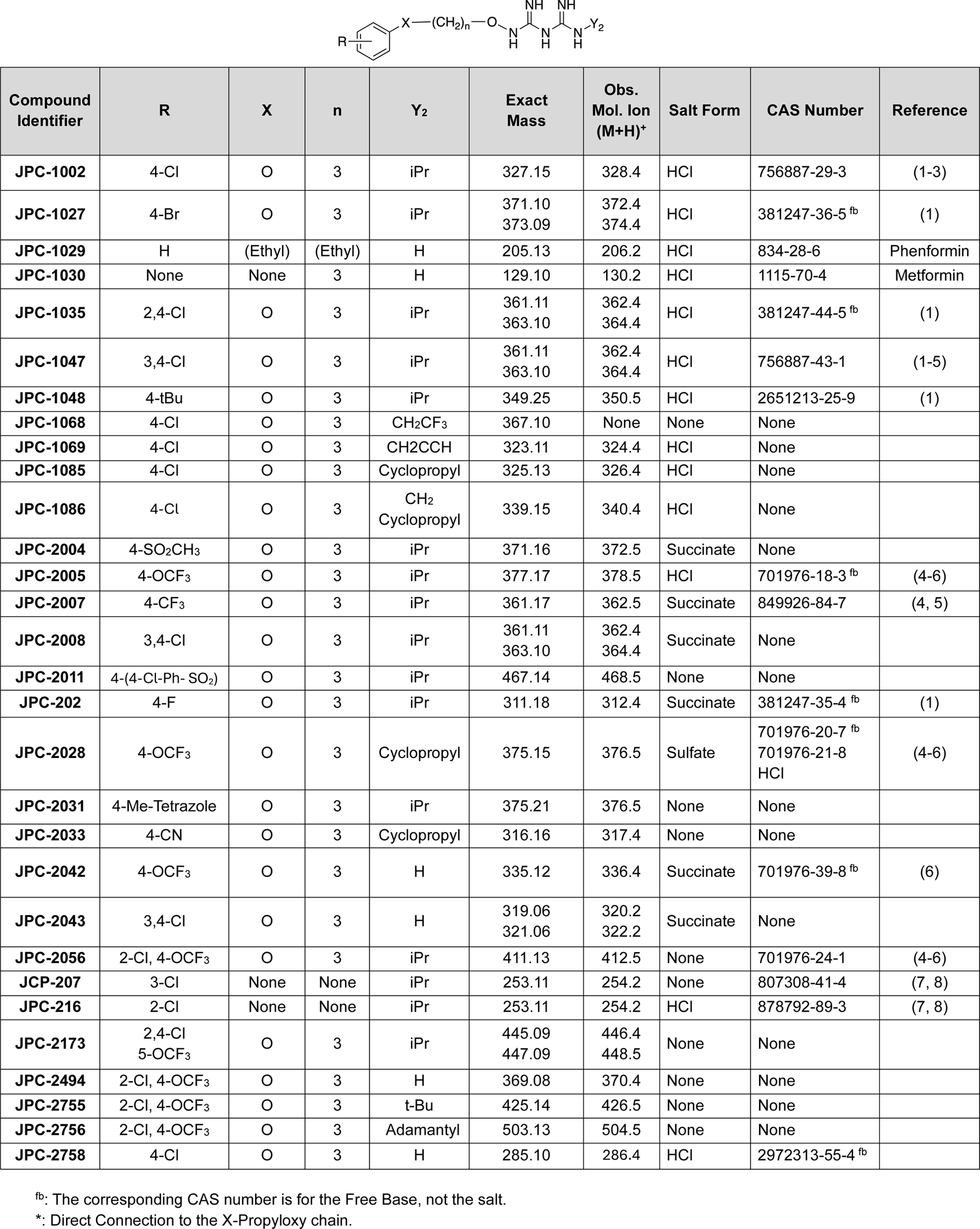
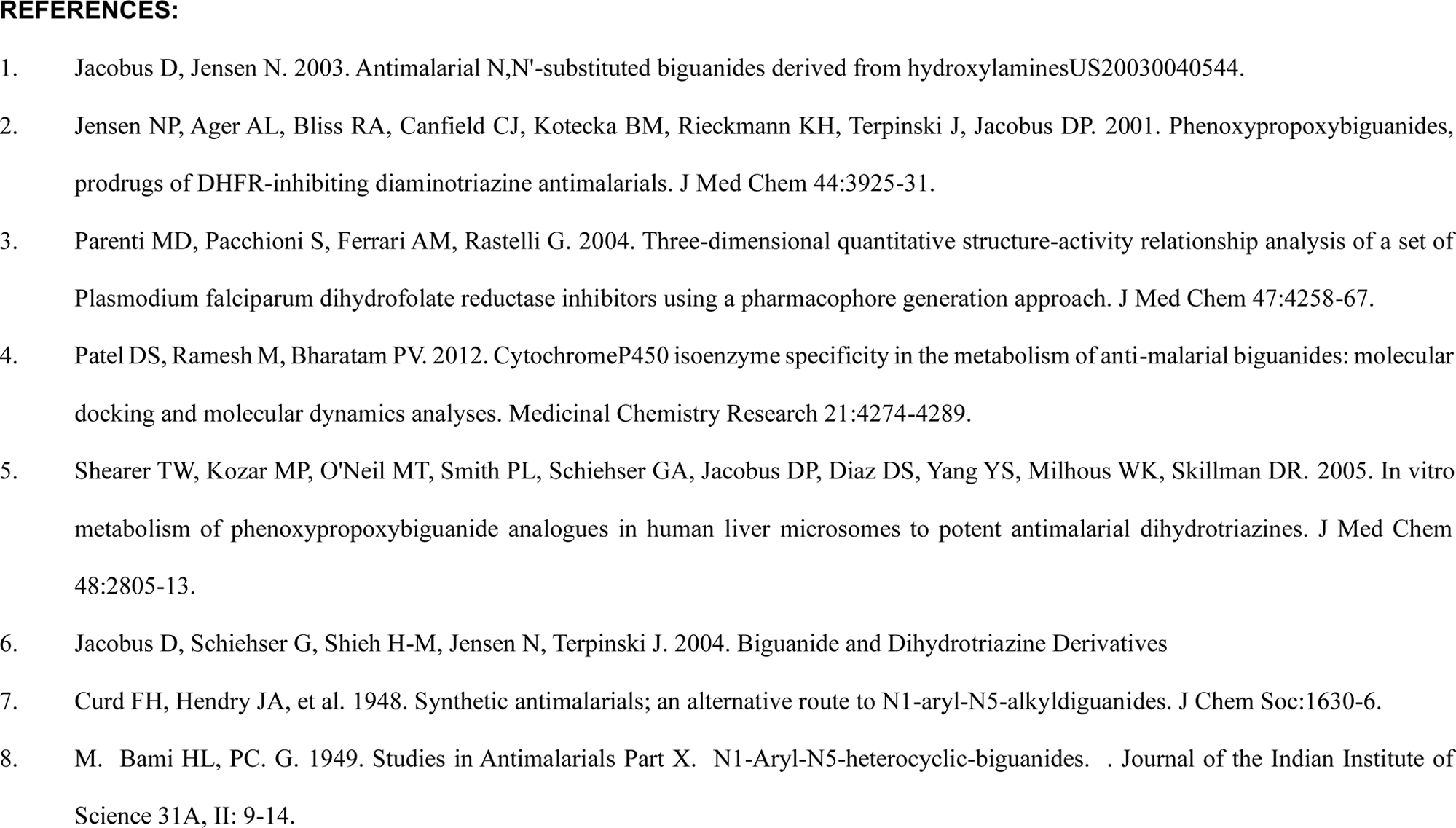
List of Biguanides with Chemical Abstract System (CAS) number, functional groups and corresponding references.

## Acknowledgments

CBM research is supported by National Institute of Health (NIH) grants (AI123321, AI138139, AI152220, and AI136118), the Steven and Alexandra Cohen Foundation (Lyme 62 2020) and the NBIA Foundations.

